# *In silico* analysis of SNPs in human phosphofructokinase, Muscle (*PFKM*) gene: An apparent therapeutic target of aerobic glycolysis and cancer

**DOI:** 10.1101/2020.05.27.118653

**Authors:** Yogita Rani, Kamaljit kaur, Madhvi Sharma, Namarta Kalia

## Abstract

Phosphofructokinase, muscle (PFKM), a key glycolytic regulatory enzyme is a potential target for cancer therapeutic studies accredited to the employed inefficient phenomenon known as Warburg effect. PFKM is encoded by *PFKM* gene located at chromosome 12q13.11. Single nucleotide polymorphisms (SNPs) are known to profoundly affect gene expression and protein function. Therefore, the first attempt was made to computationally identify putative functional PFKM variants. These SNPs were further explored to find their probable association with different cancer types. A total of 9694 SNPs were retrieved from dbSNP database. Of which, only 85 validated SNPs with ≥10% minor allele frequency (MAF) were subjected to analysis by softwares including Ensembl Genome browser, FuncPred (SNPinfo), regulomeDB (v 2.0), SIFT and PolyPhen-2. The relative analysis of output obtained classified the selected-SNPs into 11 highly prioritized (HP), 20 moderately prioritized and 54 not/poorly prioritized SNPs. The 11 HP-SNPs were found to have the highest likelihood of being functionally important, evidenced by previous association of rs2269935, rs11168417, rs11609399 and rs2228500 HP-SNPs with cachexia, lung and breast cancer. The study warrants further experiments to confirm the predictive role of prioritized SNPs in cancer etiology and also provides directions to fellow researchers.

## INTRODUCTION

Human cancers involve uncontrollable growth of abnormal cells that divides rapidly and have potential to deteriorate other body tissues. More than 100 types of cancers were reported in humans. The data interpreted by world health organization accounts for more than 9.6 million people that died globally owing to different type of cancers [1]. Beside this, 17 million new cases of people suffering from cancer were reported worldwide in 2018, which provides the estimation that the number of cancer patients would possibly increase by 2040. Different types of cancers are reported worldwide involving brain tumor, breast cancer, cervical cancer, colon cancer, lung cancer, prostate cancer *etc*. Amongst this, breast cancer is considered as most prevalent type of cancer in mammals [2]. It may affect both men and women but higher number of cases was reported in females. Various studies in oncology recommended the fact that growth of cancer cells and their metabolism is directly dependent on genetic variations [3–6].

In oncology, the phenomenon of cancer cells’ metabolism was earlier explained by Germen scientist Otto Warburg in 1920 in terms of aerobic glycolysis [7]. The metabolism of cancer cells was quite different from normal body cells. Normal body cells in humans require enough amount of oxygen for respiration and various cellular activities. Usually under aerobic conditions, cells follows oxidative phosphorylation pathway for energy production as it act as major source for ATP production. Nevertheless, under anaerobic conditions, normal cells undergoes glycolytic pathway that leads to lower energy production. This metabolic pathway involves breakdown of glucose into pyruvate molecules with an array of enzymatic reactions employing three main regulatory enzymes including *Hexokinase, Phosphofructokinase 1* (PFK1) and *Pyruvate kinase*. Both these pathways are essential for cellular metabolism as well as growth. On the contrary, even in aerobic conditions, cancer cells predominantly follows glycolytic pathway culminating in pyruvate production. This phenomenon of cancer cell respiration *via* glycolysis in aerobic conditions is designated as ‘Warburg effect’, which is a hallmark of cancer in humans [8]. Thus, this metabolism of cancer cells causes more consumption of energy (normal cells^×^10) and less ATP production, consequently leading to damage of body tissues and cellular apoptosis. Therefore, study of cancer cell metabolism and Warburg effect provides hypothesis that glycolysis plays an essential role in dissemination of human cancers [9–12].

Of the three main regulatory enzymes of glycolysis, PFK1 is the main rate limiting enzyme of glycolysis as it controls maximum percentage of glycolytic activity. PFK1 is a tetrameric protein that is allosteric in nature with enzyme activity is controlled by many activators, inhibitors and metabolites [13]. The main role of PFK1 in glycolysis is conversion of fructose-6-phosphate to fructose-1, 6-bisphosphate with the release of energy. Mammalian PFK1 spans about 30 kb and contains 24 exons, having molecular weight of 340 kDa [14–16]. Primarily, three isozymes of PFK1 have been identified in humans that include PFK-muscle (PFKM), PFK-Liver and PFK-Platelet, which function as subunits of tetrameric PFK. The molecular weight of PFKM is 85 kDa, PFKL is 80 kDa and PFKP is 85 kDa, each encoded by separate gene [17].

Of the three isozymes, a genome-wide association study has reported *PFKM* as a novel marker accountable for breast cancer in humans [18].The PFKM is encoded by *PFKM* gene, located at chromosome 12q13.11 (NCBI reference sequence number NC_000012.12, **Figure 1**). *PFKM* composed of 41,151 bases spanning the region between 48,105,253 to 48,146,404 base pairs (bp) of chromosome 12. The coding region of *PFKM* consists of 2340bp, which encodes approximately 780 amino acids [17]. Studies have explained both direct and indirect association of *PFKM* genetic mutations with different types of cancers in humans, for instance breast cancer, bladder cancer, non-small cell lung cancer, human glioma, human glioblastoma and human melanomas [3–6, 18–19]. The most common type of genetic variation investigated in these studies is single nucleotide polymorphisms (SNP).

**Figure 1:**
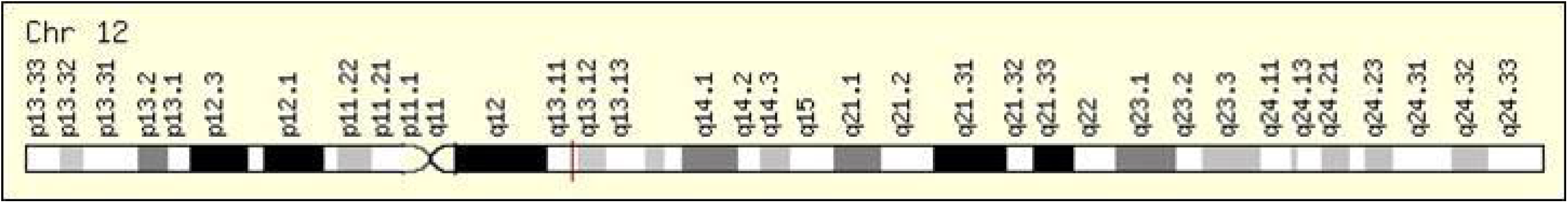
Cytogenetic Location: 12q13.11, which is the long (q) arm of chromosome 12 at position 13.11 Molecular Location: base pairs 48,105,253 to 48,146,404 on chromosome 12 (Homo sapiens Updated Annotation Release 109.20200228, GRCh38.p13) (NCBI)

Human *PFKM* consists of both coding as well as non-coding region that are responsible for total 9694 SNPs *as per* NCBI dbSNP database (dated October 2019). However, it is unlikely of all SNPs being functional (+/-) enough to be investigated in detail, therefore bioinformatics tools have been developed that can help us foretell SNPs with biological functions such as splicing, transcriptional, translational, miRNA binding, protein stability *Electra* [20]. Additionally, there are some other factors such as significance of function identified, validation status, presence of SNPs in evolutionary conserved region, minor allele frequency (MAF) that can further support SNP prioritization [21]. Thus, the present study is based on scrutinizing the functionally important SNPs of *PFKM* from the list of casual variants and finding their probable association with cancer using *in silico* analysis and literary evidences. The results of the research are predicted after comparison of output provided by different bioinformatics tools used in the present study. As per our finest knowledge, this is the first computational analysis performed on human *PFKM* to identify functionally important casual variants of the human *PFKM*, which might be related to cancer proliferation in humans.

## METHODS

*In silico* analysis was performed to identify the functional SNPs of *PFKM* present in human muscles at chromosome 12q13.11 (NCBI). SNPs that represent any functional and structural impacts were selected on the basis of methods adapted in our research for computational analysis of human *PFKM* as illustrated in **Figure 2**.

**Figure 2:**
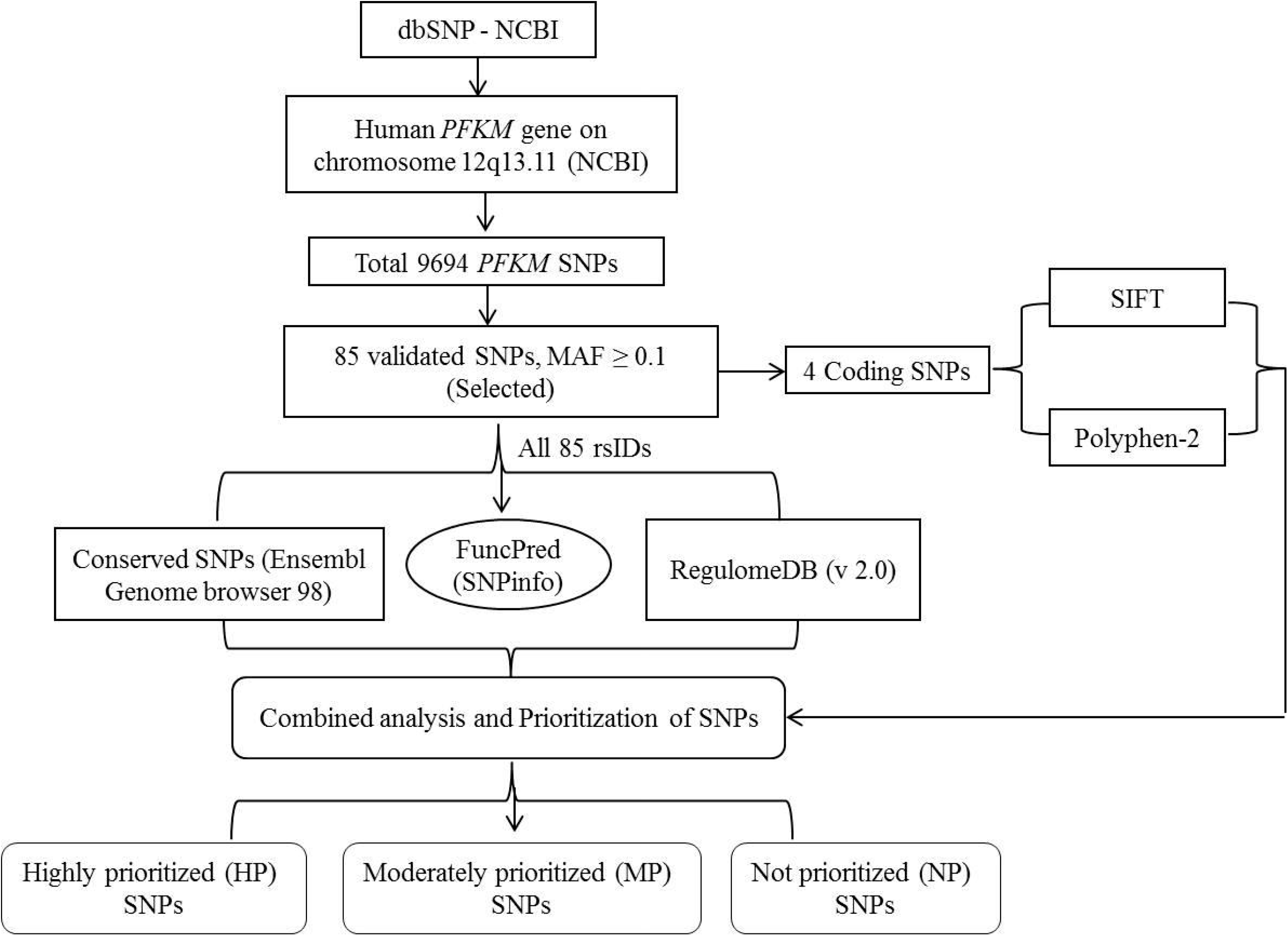
Schematic representation of computational analysis for scrutinizing validated functional SNPs of human *PFKM*.

### Recruiting PFKM polymorphisms from dbSNP database

The data of putatively functional SNPs for *PFKM* (*Homo sapiens*) was obtained from dbSNP (https://www.ncbi.nlm.nih.gov/snp), which is most extensive database of genomic variations [22]. SNP gene view showed the presence of total 9694 SNPs of *PFKM* in gene region (**S. Fig. 1**). From this aggregate, four parameters were used to prioritize SNPs that includes confirmation of validation status, presence in evolutionary conserved region, MAF ≥ 0.10 and significance of functions identified. Thus, those SNPs that fulfilled the above criteria were selected and further explored in the present study.

### Identifying evolutionary conserved SNPs of PFKM

Manual spotting of *PFKM* SNPs was performed in the evolutionary conserved regions using the Ensembl Genome browser 98 (https://asia.ensembl.org/index.html). For this, 91 eutherian mammals showing genomic alignments were comparatively analyzed with the human *PFKM*, which involves various species including Drill, Pig-tailed macaque, Angola colobus, Bolivian squirrel monkey, Mouse lemur, American beaver, Kangaroo rat, Squirrel, Chinese hamster PICR, Golden hamster, Tarsier, Bushbaby, Greater bamboo lemur, Damara mole rat, Naked mole-rat male, Naked mole-rat female, Mongolian gerbil, Gibbon, Chimpanzee, Gorilla, Olive baboon, Gelada, Vervet-AGM, Capuchin, Degu *etc*. (**S. Fig. 2**).

The base pair view from database was preferred for manual verification of SNPs and detailed comparison of alignments of various species with human *PFKM* was performed. Ensembl genome browser 98 helps in investigating conserved SNPs (cSNPs) and provides information regarding polymorphisms in gene sequences. Moreover, genomic alignments also help to differentiate between the conserved and non-conserved regions in genome. The data available on the browser is in FASTA format can also be retrieved whenever needed for further study. The highlighted regions (yellow, green, purple, pink and red) with hyperlinks in the region of sequence alignments represents the variant SNPs, which can be studied in detail, by clicking them a dialogue box that appeared on the screen as depicted in **Figure 3**.

**Figure 3:**
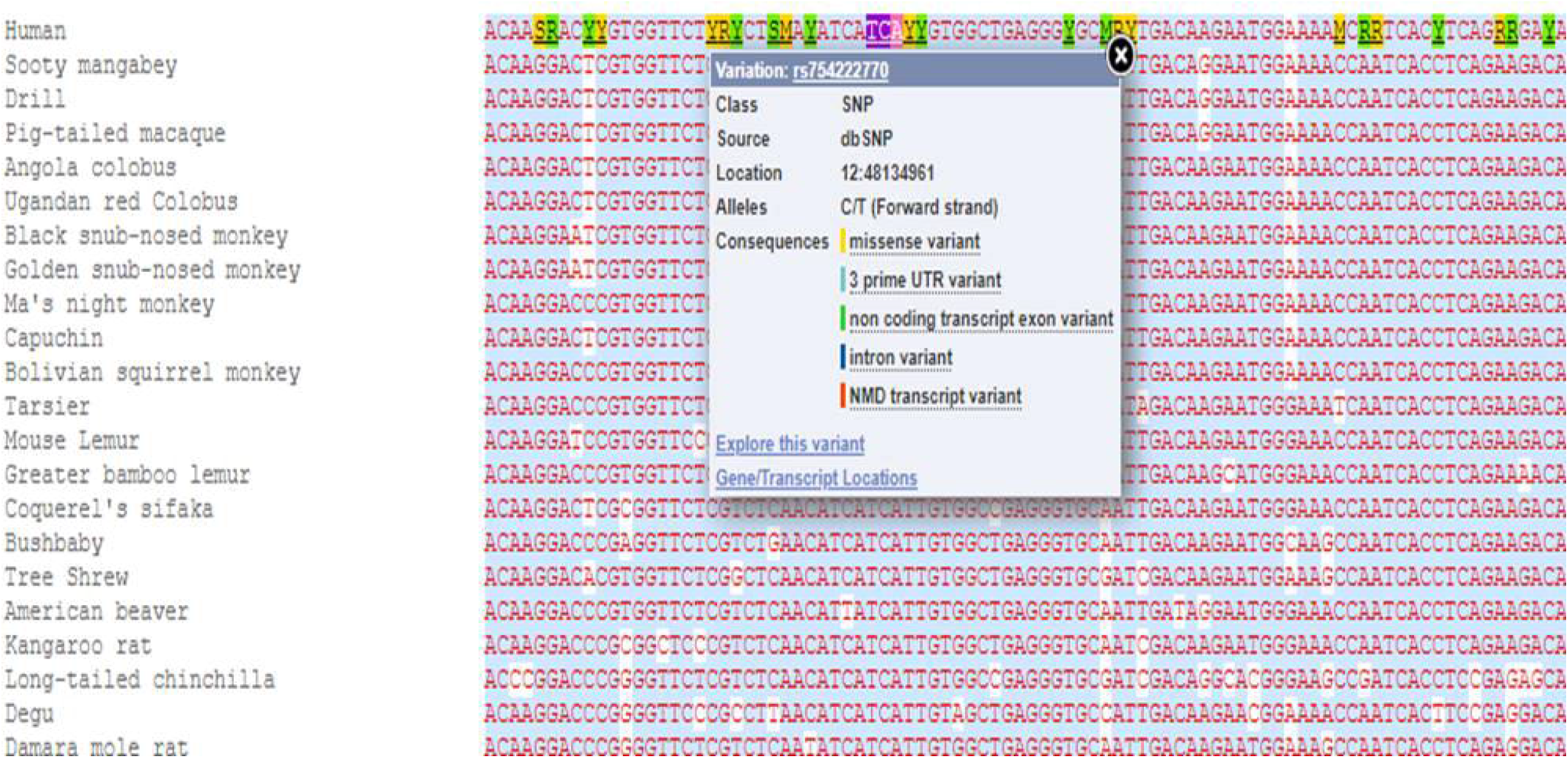
The image showing comparison of the human *PFKM* gene with some species from 91 Eutherian mammals; Human *PFKM* gene sequences indicated in red color, Conserved region represented by blue highlighted region. The highlighted (yellow, green, purple) nucleotides indicate variant SNPs. This screenshot was taken from Ensembl Genome browser 98.

### Recognition of likely functional PFKM variants

Identification of functional effects of *PFKM* SNPs was performed using two standard bioinformatics tools *i.e*. SNPinfo (FuncPred) and RegulomeDB (version 2.0). These tools predicts SNPs with potential functions such as transcription factor binding sites (TFBS), Intron/exon border consensus sequences (splice sites), exonic splicing enhancers (ESEs) and miRNA binding.

#### 1. SNP analysis using SNPinfo (FuncPred)

SNPinfo (FuncPred) (https://snpinfo.niehs.nih.gov/snpinfo/snpfunc.html) provides collection of functional information associated with query SNPs of different genes using variety of resources and algorithms [23]. In our research, this web server was used in Asian population (ASW). The investigators can query finite list of SNPs for a particular set of gene of interest for identifying SNPs with potential functions. This web browser also helps in the selection of SNPs for genetic association studies. It consists of a composite tool for SNP function prediction thus named as ‘FuncPred’. For the present study, a list of rsIDs of validated SNPs with MAF ≥0.10 was uploaded in FuncPred for batch analysis with default settings in Asian population as shown in **S. Fig. 3**. The output obtained represents the list of putatively functional SNPs of human *PFKM*.

#### 2. SNP analysis using RegulomeDB (v 2.0)

To predict the role of various polymorphisms in a specific gene Beta Regulome or RegulomeDB (v 2.0) is mostly availed. This database describes the transcriptional role of variants as regulatory elements of gene [24]. For this SNPs were analyzed using RegulomeDB (v 2.0) (https://beta.regulomedb.org/regulome-search/); an online composite bioinformatics database [25]. The database include high quality updated datasets from Encyclopedia of DNA Elements transcription factor, chromatin immunoprecipitation sequencing, DNAse I hypersensitive site data and many other sources like dsQTL, ChIP-exo. A list of dbSNP rsIDs was used as an input for analysis by software (**S. Fig. 4**). The annotation scores to the variants were assigned according to the information available in the database that classifies the variants mainly into six categories ranging from 1 to 6 as per scoring scheme.

Category 7 has been also assigned to some variants which represents that no current data availability about these variants but it may be possible that these variants may show functional effects in future. The category 1 variants represents that they were ‘likely to affect binding and linked to target gene expression’, category 2 variants were ‘likely to affectbinding, category 3 variants were ‘less likely to affect binding’. Moreover, category 4, 5 and 6 variants represents the ‘minimal binding evidence’. The annotation scores assigned to variants may be subdivided into sub-categories on the basis of their behavior of binding and functional effects.The scoring was based on the data available database but it was not considered as final information of variants; because it may varies alongwith apprising of database; as per further researches carried out in future. The data provided in **S. Table 1** represents the scheme of RDB scoring [24] which was used in our research for computational analysis of functional SNPs of *PFKM*.

### Functional effect prediction of the translated SNPs of PFKM

The damaging and deleterious effects of SNPs of coding region of gene was predicted using *SIFT* and *Polyphen-2*. These are basic computational tools with high concordance of testing human genes’ mutations and their lethal effects on structure of proteins and genes [26]. These tools provide information of genetic variants and thus help in prioritization of variants with functional characters. The variants lying in coding region were tested for their concordance, sensitivity and specificity, data provided in (+/-) predictive values [27]. These two softwares similarly predict pathogenicity of gene of interest [28].

#### 1. SNP analysis using SIFT(Sorting Intolerant From Tolerant)

SIFT (https://sift.bii.a-star.edu.sg/) is a web based computational tool used for analyzing coding SNPs of human *PFKM* using different algorithms. Interpretation by this tool was expressed in the form of table classifying the effect of mutation of SNP listed along with SIFT score, median and prediction details. The rsID of SNP was uploaded to get possible predictions. Predictions were generated with default settings of SIFT program such as SNP may be designated either as ‘Tolerated’ or ‘Not Tolerated’ [29]. The sensitivity and specificity of any variant can be determined by the percentage and scoring (0–1) predicted by program. Scores lying within range (0.00-0.005) consider variant as ‘damaging’, score (0.0051-0.10) considered as ‘potentially damaging’, score (0.101-0.20) considered as ‘borderline’ and score (0.201-1.00) was considered as ‘tolerant’ [21].

#### 2. SNP analysis using Polyphen-2 (Polymorphism Phenotyping v-2)

Polyphen-2 (http://genetics.bwh.harvard.edu/pph2/) is the standard tool which is sequenced based, used for coding variants analysis. It estimates protein sequence variation and protein function [30]. The algorithm of tool relies on both physical and comparative considerations to predict possible functional consequences of amino acid substitutions on gene structure and function. This tool calculates the position specific independent count (PSIC) score for genetic variants. The scoring provided by this tool (0 – 1) represents the probability of damaging effects of variant depending upon the threshold value. If the score predicted by the tool lies above the threshold value (score > 0.2) then effect is considered as ‘probably damaging’ and ‘possibly damaging’ (score > 0.2 and < 0.96) but if score lies below threshold value the effect is said to be ‘Benign’ (score < 0.2) or ‘Neutral’ [27]. The possible effects of variants are predicted by inputting rsIDs of SNPs along with amino acid substitution position details. A vertically marked colored gradient bar is shown as result by Polyphen-2 for the variant, the green region in the bar represents effect to be ‘Benign’ and red region represents the ‘Damaging’ effect alongwith Polyphen-2 score.

## RESULTS

Human *PFKM* consists of total 9694 SNPs in dbSNP database. From these 9694 SNPs, 85 SNPs were found to be validated and further had MAF ≥ 0.10 were filtered out for subsequent *in silico* analysis (**Table 1**). Other SNPs were excluded from the present analysis as they neither had MAF ≥ 0.10 nor they were found to be validated. These 85 SNPs were spanning both coding as well as non-coding regions. For instance, two SNPs were of non-coding region particularly the mis-sense or non-synonyms mutations (*ns-SNPs*; rs11609399 and rs2228500), two SNPs were synonymous SNPs (*syn-SNPs*; rs1049392 and rs8716), four SNPs identified of 5’ near gene region (rs146586156, rs11168408, rs10875743 and rs10875744) while, rest of the 77 SNPs were identified as intronic. Thus, our investigation accounted for 85 SNPs of human *PFKM* belonging to different regions. The *PFKM* sequence of 91 eutherian mammals showed alignment with human *PFKM* in Ensembl genome browser 98, which depicted only four SNPs conserved among these species in spite of evolutionary divergence, hence, referred as conserved SNPs (cSNPs) in the present study. These cSNPs includes rs11609399, rs2228500, rs1049392 and rs8716.

**Table 1 –.** List of 85 SNPs (validated with MAF ≥ 0.10) source dbSNP database.

| S.No. | SNP rsID | SNPs' Types | Chromosome Position | Heterozygosity | Validation Status | MAF |
| --- | --- | --- | --- | --- | --- | --- |
| 1. | rs146586156 | 5' near | 48104312 | 0.453 | 1 2 6 | 0.3462 |
| 2. | rs11168408 | 5' near | 48104544 | 0.295 | 1 2 4 5 6 | 0.1799 |
| 3. | rs10875743 | 5' near | 48104630 | 0.329 | 1 2 4 5 6 | 0.2077 |
| 4. | rs10875744 | 5' near | 48104657 | 0.478 | 1 2 4 5 6 | 0.3954 |
| 5. | rs4760619 | intron | 48106148 | 0.188 | 1 2 6 | 0.1036 |
| 6. | rs726354 | intron | 48106551 | 0.459 | 1 2 3 4 6 | 0.3562 |
| 7. | rs534307036 | intron | 48106710 | 0.395 | 1 2 6 | 0.2708 |
| 8. | rs11609399 | Mis-sense | 48107378 | 0.391 | 1 2 4 5 6 | 0.3562 |
|  | □ |  |  |  |  |  |
| 9. | rs10492080 | intron | 48107628 | 0.459 | 1 2 4 5 6 | 0.3562 |
| 10. | rs725454 | intron | 48108317 | 0.272 | 1 2 4 6 | 0.1621 |
| 11. | rs757556 | intron | 48108710 | 0.459 | 1 2 4 5 6 | 0.3562 |
| 12. | rs17122666 | intron | 48108791 | 0.251 | 1 2 6 | 0.1474 |
| 13. | rs12099462 | intron | 48109321 | 0.330 | 1 2 4 6 | 0.2087 |
| 14. | rs879242 | intron | 48109518 | 0.459 | 1 2 4 6 | 0.3562 |
| 15. | rs72644845 | intron | 48110098 | 0.459 | 1 2 6 | 0.3562 |
| 16. | rs4760681 | intron | 48111447 | 0.273 | 1 2 4 6 | 0.1633 |
| 17. | rs58331734 | intron | 48111609 | 0.459 | 1 2 6 | 0.3564 |
| 18. | rs11613781 | intron | 48113122 | 0.459 | 1 2 4 5 6 | 0.3562 |
| 19. | rs57655233 | intron | 48113603 | 0.459 | 1 2 6 | 0.3562 |
| 20. | rs2158515 | intron | 48113885 | 0.478 | 1 2 4 6 | 0.3954 |
| 21. | rs7302354 | intron | 48114047 | 0.305 | 1 2 4 5 6 | 0.1875 |
| 22. | rs2107638 | intron | 48114136 | 0.309 | 1 2 4 6 | 0.1909 |
| 23. | rs78649179 | intron | 48114413 | 0.459 | 1 2 6 | 0.3564 |
| 24. | rs72644846 | intron | 48114500 | 0.459 | 1 2 6 | 0.3562 |
| 25. | rs11168412 | intron | 48114703 | 0.229 | 1 2 6 | 0.1318 |
| 26. | rs61343568 | intron | 48114863 | 0.496 | 1 2 6 | 0.4565 |
| 27. | rs60106007 | intron | 48115981 | 0.373 | 1 2 6 | 0.2482 |
| 28. | rs12228924 | intron | 48115987 | 0.304 | 1 2 6 | 0.1873 |
| 29. | rs76341202 | intron | 48116137 | 0.283 | 1 2 6 | 0.1707 |
| 30. | rs113785514 | intron | 48117174 | 0.373 | 1 2 6 | 0.2482 |
| 31. | rs11168413 | intron | 48117212 | 0.309 | 1 2 4 6 | 0.1909 |
| 32. | rs11168414 | intron | 48118548 | 0.297 | 1 2 6 | 0.1484 |
| 33. | rs11168415 | intron | 48118738 | 0.373 | 1 2 6 | 0.2484 |
| 34. | rs5798062 | intron | 48118890 | 0.296 | 1 2 6 | 0.1807 |
| 35. | rs12306290 | intron | 48119303 | 0.373 | 1 2 4 6 | 0.2480 |
| 36. | rs11168416 | intron | 48120196 | 0.373 | 1 2 4 6 | 0.2478 |
| 37. | rs10875745 | intron | 48121622 | 0.308 | 1 2 4 6 | 0.1901 |
| 38. | rs11168417 | intron | 48121937 | 0.370 | 1 2 4 6 | 0.2446 |
| 39. | rs11168418 | intron | 48121971 | 0.273 | 1 2 6 | 0.1629 |
| 40. | rs11168419 | intron | 48121980 | 0.370 | 1 2 4 6 | 0.2446 |
| 41. | rs11168420 | intron | 48122113 | 0.292 | 1 2 6 | 0.1777 |
| 42. | rs72644847 | intron | 48122310 | 0.425 | 1 2 6 | 0.3059 |
| 43. | rs4760683 | intron | 48122513 | 0.483 | 1 2 4 6 | 0.4077 |
| 44. | rs2269935 | intron | 48122681 | 0.241 | 1 2 6 | 0.1402 |
| 45. | rs57642441 | intron | 48123524 | 0.370 | 1 2 6 | 0.2446 |
| 46. | rs10082861 | intron | 48123615 | 0.370 | 1 2 4 6 | 0.2448 |
| 47. | rs4760684 | intron | 48124297 | 0.370 | 1 2 4 6 | 0.2452 |
| 48. | rs10875746 | intron | 48124481 | 0.301 | 1 2 4 6 | 0.1849 |
| 49. | rs11168421 | intron | 48124550 | 0.368 | 1 2 4 6 | 0.2430 |
| 50. | rs73104071 | intron | 48124786 | 0.307 | 1 2 6 | 0.1893 |
| 51. | rs58540891 | intron | 48124796 | 0.307 | 1 2 6 | 0.1893 |
| 52. | rs1859445 | intron | 48124808 | 0.313 | 1 2 3 4 6 | 0.1943 |
| 53. | rs562841214 | intron | 48125621 | 0.484 | 1 2 6 | 0.4107 |
| 54. | rs11168422 | intron | 48125722 | 0.278 | 1 2 6 | 0.1671 |
| 55. | rs4760685 | intron | 48127186 | 0.491 | 1 2 4 5 6 | 0.4333 |
| 56. | rs11374838 | intron | 48127237 | 0.252 | 1 2 6 | 0.1480 |
| 57. | rs527761146 | intron | 48128263 | 0.452 | 1 2 6 | 0.3317 |
| 58. | rs4141065 | intron | 48129716 | 0.458 | 1 2 6 | 0.3548 |
| 59. | rs1476607 | intron | 48131021 | 0.297 | 1 2 3 4 6 | 0.1811 |
| 60. | rs10875747 | intron | 48132078 | 0.295 | 1 2 4 6 | 0.1546 |
| 61. | rs10875748 | intron | 48132577 | 0.261 | 1 2 4 6 | 0.1546 |
| 62. | rs2228500 □ | Mis-sense | 48132929 | 0.292 | 1 2 | 0.1478 |
| 63. | rs2269933 | intron | 48133276 | 0.387 | 1 2 4 6 | 0.2444 |
| 64. | rs1049392 □ | syn-SNP | 48133403 | 0.253 | 1 2 6 | 0.1879 |
| 65. | rs11168424 | intron | 48133932 | 0.307 | 1 2 6 | 0.1893 |
| 66. | rs2286021 | intron | 48134437 | 0.252 | 1 2 6 | 0.1480 |
| 67. | rs2286020 | intron | 48134636 | 0.307 | 1 2 3 6 | 0.1891 |
| 68. | rs10219623 | intron | 48135766 | 0.313 | 1 2 6 | 0.1943 |
| 69. | rs80140486 | intron | 48135913 | 0.311 | 1 2 6 | 0.1923 |
| 70. | rs78294086 | intron | 48135915 | 0.308 | 1 2 6 | 0.1899 |
| 71. | rs77966213 | intron | 48135916 | 0.307 | 1 6 | 0.1895 |
| 72. | rs75261947 | intron | 48135920 | 0.253 | 1 6 | 0.1484 |
| 73. | rs79336782 | intron | 48135919 | 0.307 | 1 6 | 0.1895 |
| 74. | rs75333732 | intron | 48135922 | 0.307 | 1 6 | 0.1895 |
| 75. | rs79751478 | intron | 48135925 | 0.297 | 1 6 | 0.1811 |
| 76. | rs76706444 | intron | 48135928 | 0.298 | 1 2 6 | 0.1821 |
| 77. | rs10219645 | intron | 48135954 | 0.368 | 1 2 4 6 | 0.2434 |
| 78. | rs10875749 | intron | 48138134 | 0.334 | 1 2 4 5 6 | 0.2119 |
| 79. | rs11168425 | intron | 48138414 | 0.369 | 1 2 6 | 0.2442 |
| 80. | rs58556309 | intron | 48140505 | 0.306 | 1 2 6 | 0.1889 |
| 81. | rs34700912 | intron | 48141031 | 0.491 | 1 2 6 | 0.4341 |
| 82. | rs148905596 | intron | 48143390 | 0.313 | 1 2 6 | 0.1939 |
| 83. | rs4075913 | intron | 48143841 | 0.389 | 1 2 4 5 6 | 0.3546 |
| 84. | rs11168427 | intron | 48145196 | 0.292 | 1 2 6 | 0.1490 |
| 85. | rs8716 □ | syn-SNP | 48145699 | 0.436 | 1 2 4 6 | 0.2043 |

MAF = Minor allele frequency $\geq 0.10$ , syn-SNP=Synonymous SNP, □ denote conserved SNPs (cSNPs), 1 to 6 numbers represents validation status as below:
1. Validated by multiple, independent submissions to the ref SNP cluster 2. Validated by frequency or genotype data: minor alleles observed in at least two chromosomes. 3. Validated by submitter confirmation 4. All alleles have been observed in at least two chromosomes apiece 5. Genotyped by HapMap project 6. SNP has been sequenced in 1000Genome project.

FuncPred analysis of 85 selected SNPs predicted total 22 SNPs with important regulatory function (**Table 2**). Amongst these 22 SNPs, 21 SNPs were found to be affecting transcription factor binding site. Moreover, two SNPs, rs11609399 and rs2228500 were predicted as exonic splicing effectors. The rest 63 SNPs from the selected SNPs list did not show any information regarding functional effects in FuncPred database. The selected 85 SNPs were also analyzed in Beta Regulome database or RegulomeDB (v 2.0) to check the annotation scores assigned by the database. The rsIDs of SNPs were incorporated as an input in the database, which divided 85 validated SNPs into six broad categories *i.e*. category 1 to category 6 [25]. It was found that from overall 85 SNPs, only 73 SNPs showed an annotation scores within category 1 to 6 (**Table 3**), while for the rest 12 SNPs (not shown in **Table 3**) the database provided no information, which is classified into category 7 *i.e*. no data availability. From the 73 identified SNPs, rs11168417 and rs2286020 were predicted as top ranked SNPs with annotation score ‘1d’ and ‘1f’ respectively. Both the scores indicate that these two SNPs have important regulatory functions for the expression of targeted gene. Further, 6 SNPs (rs11609399, rs4760619, rs5798062, rs12306290, rs11168416, rs57642441) had annotation score of ‘2b’ (likely to affect binding), 10 SNPs showed annotation score 3a (less likely to affect binding), 55 SNPs showed ‘minimum functional evidence’with category 4, 5 and 6 (**Table 3**).

**Table 2-.**
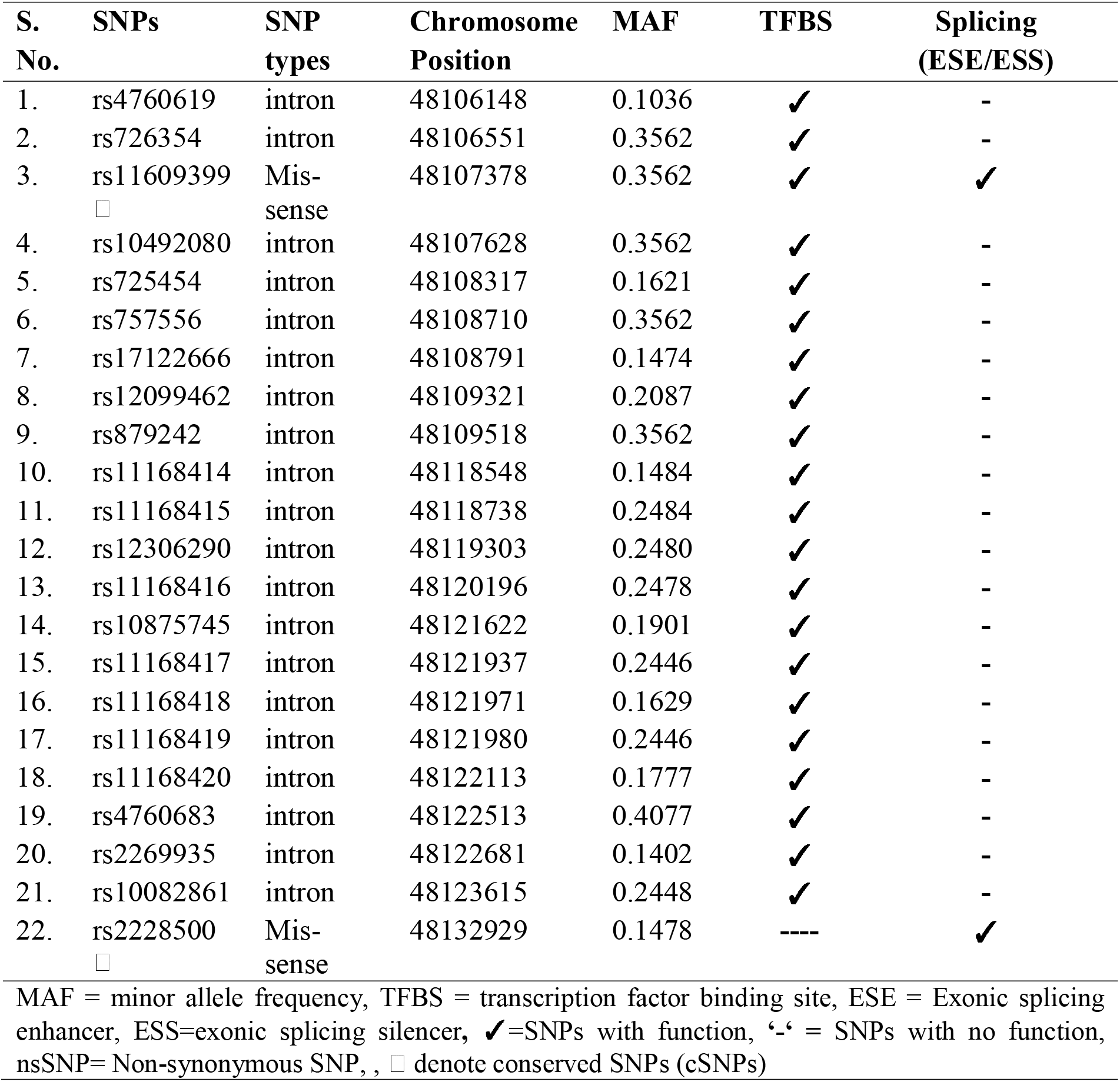
List of validated SNPs predicted by SNPinfo (FuncPred) as functional

**Table 3-.**
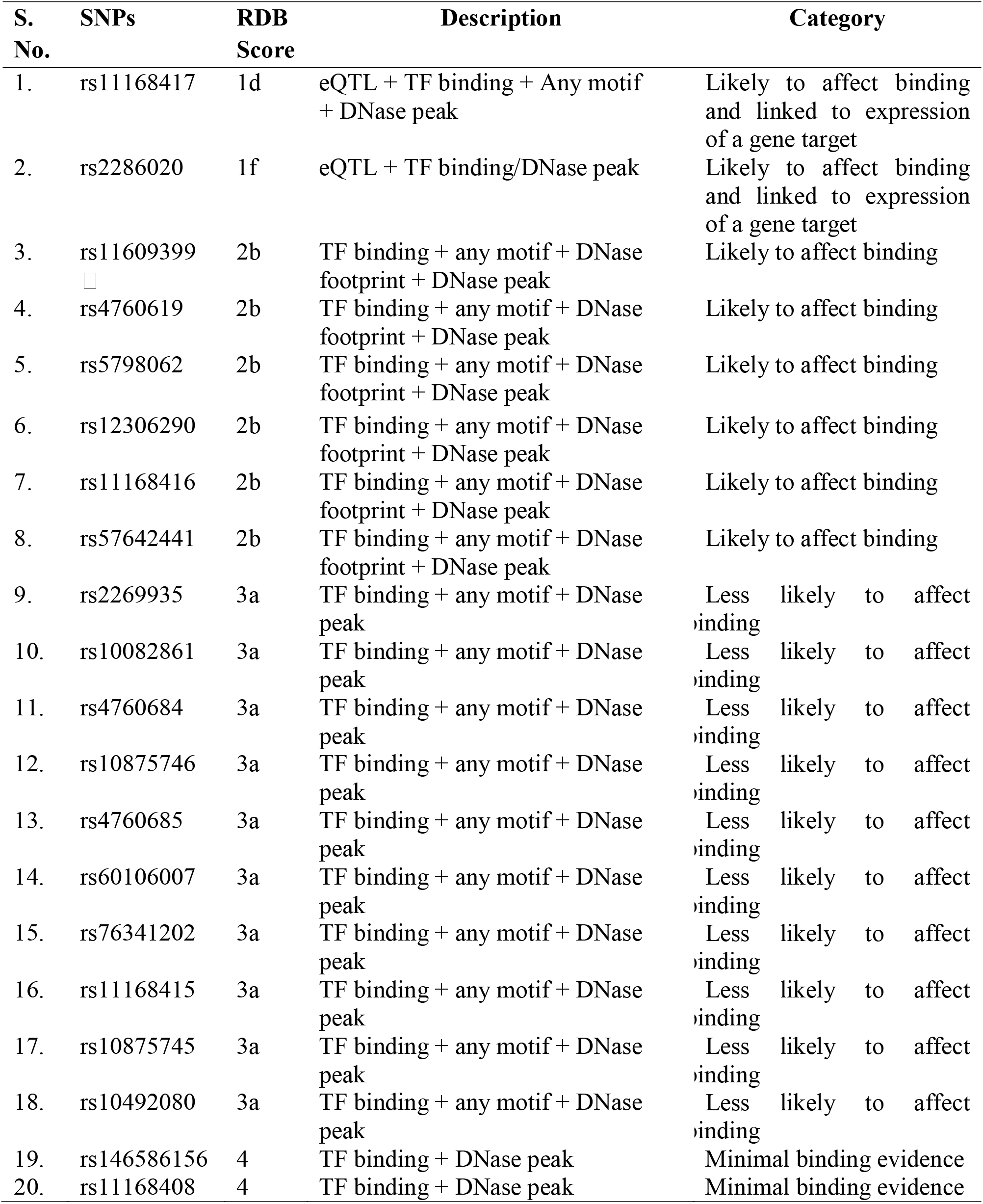
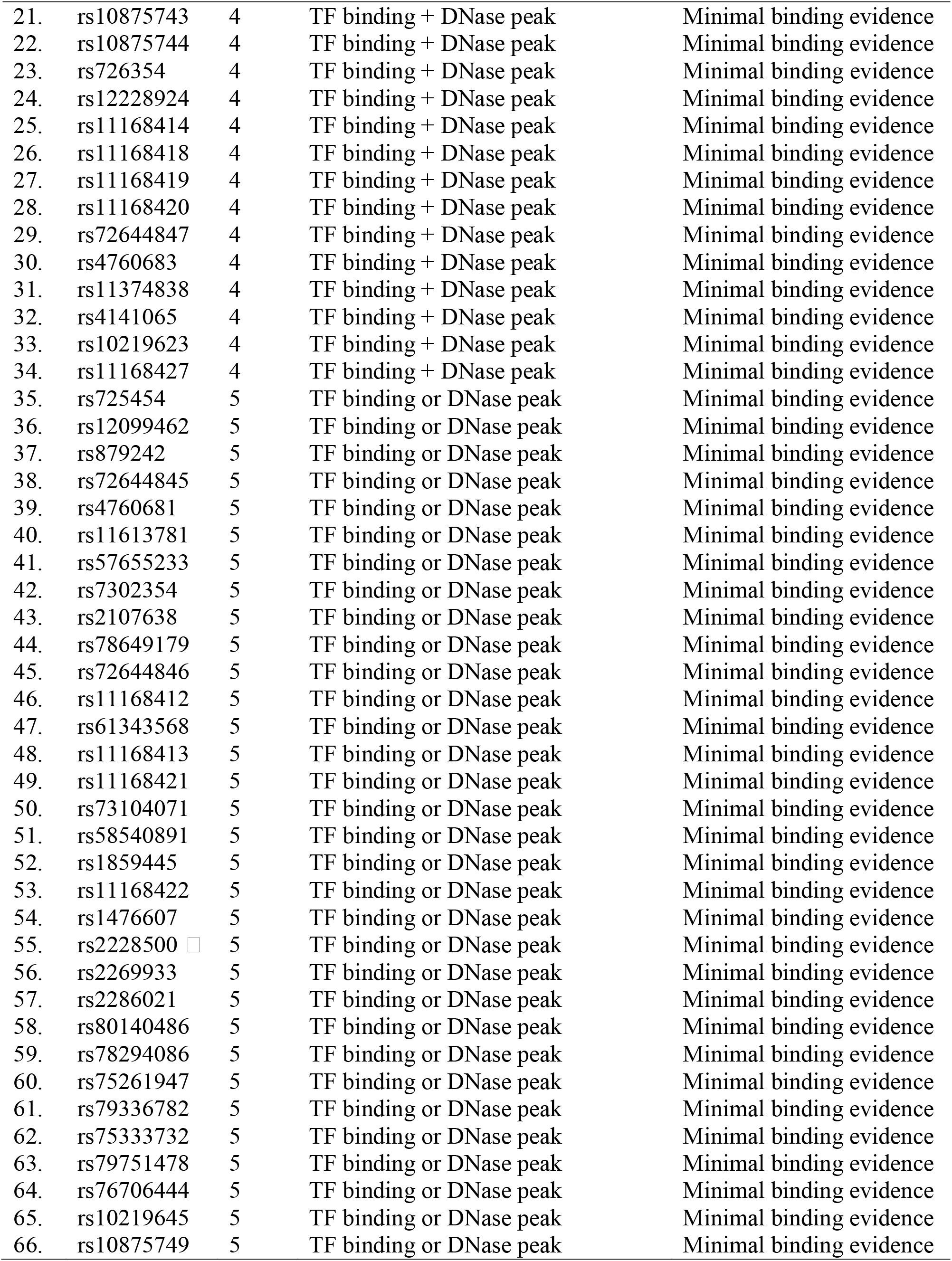
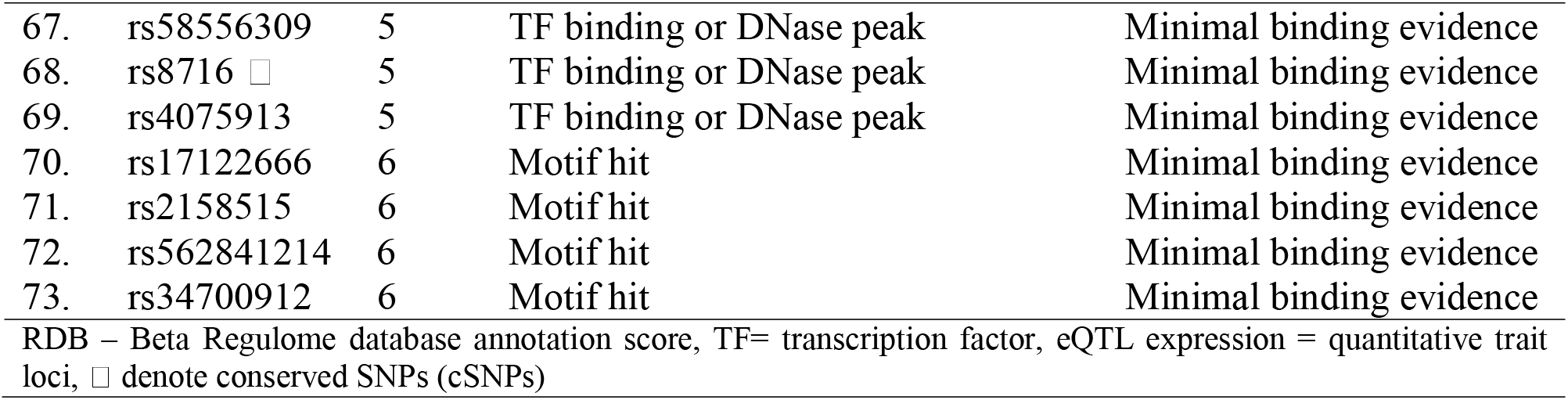
Validated SNPs listed based on the score provided by RegulomeDB (v 2.0)

Of the 85 select SNPs, four SNPs lies in exonic region, with rs11609399 and rs2228500 were ns-SNPs, while rs1049392 and rs8716 were syn-SNPs. These SNPs were analyzed using SIFT and Polyphen-2 to predict the damaging effect of these mutations on PFKM (**Table 4**). It is well-known that syn-SNPs do not affect the morphology of encoded protein as it codes for same amino-acid, therefore the chances of syn-SNPs to be found damaging was null and the same was interpreted by the softwares. The ns-SNP rs11609399 was found to be putatively deleterious with low confidence SIFT score (**S. Fig. 5**). However, Polyphen-2 did not show any information for this SNP and showed no data availability (**Table 4**). Furthermore, both SIFT and Polyphen-2 predicted ns-SNP rs2228500 as non-deleterious SNP (**S. Fig. 6**). The colored gradient vertical bar predicted by Polyphen-2 represents the variant threshold lying within green zone thus suggesting non-damaging effect of the SNP on protein structure.

**Table 4–.** Predictions of analysis of translated SNPs of human *PFKM* in *Polyphen-2* and *SIFT*

| S. No. | SNPs | Amino-acid substitution and position (NCBI) | *Polyphen-2* | *SIFT* |  |  |
| --- | --- | --- | --- | --- | --- | --- |
|  |  |  |  | SIFT Score | SIFT Median | SIFT Prediction |
| 1. | rs11609399 | Leucine to Histidine at position '2' | Data not available | 0 | 4.32 | Deleterious |
| 2. | rs2228500 | Glutamine to Arginine at position '171' | Benign Score = 0.003 | 0.352 | 2.55 | Tolerated |
| 3. | rs1049392 □ | Threonine to Threonine at position '243' | Benign Score = 0.000 | 1 | 2.56 | Tolerated |
| 4. | rs8716 □ | Alanine to Alanine at position '849' | Benign Score = 0.000 | 1 | 4.32 | Tolerated |

The comparative analysis of results predicted by different algorithms was carried out to prioritize important SNPs that are probable to be more functional than others (**Table 5**). FuncPred predictions (*i.e*. transcription factor binding site or splicing effectors), RDB annotation score from 1 to 3, cSNPs, deleterious and damaging prediction of *SIFT* and *Polyphen-2* were interpreted as functional. Based on this interpretation, the 85 SNPs were ordered from highly prioritized (HP) to Not/poorly prioritized (NP) SNPs. Those SNPs that were identified to be putatively functional by two or more tools were categorized as HP including rs11609399, rs11168417, rs4760619, rs12306290, rs11168416, rs10492080, rs11168415, rs10875745, rs2269935, rs10082861and rs2228500 (**Table 5**). These 11 HP-SNPs have higher odds of being functional than other SNPs of *PFKM*. Those SNPs that were identified to be putatively functional by one tool only were categorized as moderately prioritized (MP). Total 19 SNPs were MP, which means these SNPs have sufficient odds of being functional enough. Finally, the 55 SNPs that were not identified by any software to have function were referred as NP depicting the negligible odds of these SNPs to be functional enough to be studied. Thus, overall 11SNPs including rs11609399, rs11168417, rs4760619, rs12306290, rs11168416, rs10492080, rs11168415, rs10875745, rs2269935, rs10082861 and rs2228500 of human *PFKM* were scrutinized through a plethora of SNPs and are predicted as casual variants to be studied in detail.

**Table 5–.** Comparative analysis of 85 selected SNPs based on the results of tools used.

| S. No. | SNP rsID | SNPs' types | FuncPred | RDB Score | SIFT | Polyphen-2 | SNP-Ranking |
| --- | --- | --- | --- | --- | --- | --- | --- |
| 1. | rs11609399<br>□ | Mis-sense | □ | 2b | Deleterious | NF | HP |
| 2. | rs11168417 | intron | □ | 1d | ---- | ---- | HP |
| 3. | rs4760619 | intron | □ | 2b | ---- | ---- | HP |
| 4. | rs12306290 | intron | □ | 2b | ---- | ---- | HP |
| 5. | rs11168416 | intron | □ | 2b | ---- | ---- | HP |
| 6. | rs10492080 | intron | □ | 3a | ---- | ---- | HP |
| 7. | rs11168415 | intron | □ | 3a | ---- | ---- | HP |
| 8. | rs10875745 | intron | □ | 3a | ---- | ---- | HP |
| 9. | rs2269935 | intron | □ | 3a | ---- | ---- | HP |
| 10. | rs10082861 | intron | □ | 3a | ---- | ---- | HP |
| 11. | rs2228500 □ | Mis-sense | □ | 5 | Tolerated | Benign | HP |
| 12. | rs2286020 | intron | ---- | 1f | ---- | ---- | MP |
| 13. | rs5798062 | intron | ---- | 2b | ---- | ---- | MP |
| 14. | rs57642441 | intron | ---- | 2b | ---- | ---- | MP |
| 15. | rs60106007 | intron | ---- | 3a | ---- | ---- | MP |
| 16. | rs76341202 | intron | ---- | 3a | ---- | ---- | MP |
| 17. | rs4760684 | intron | ---- | 3a | ---- | ---- | MP |
| 18. | rs10875746 | intron | ---- | 3a | ---- | ---- | MP |
| 19. | rs4760685 | intron | ---- | 3a | ---- | ---- | MP |
| 20. | rs726354 | intron | □ | 4 | ---- | ---- | MP |
| 21. | rs11168414 | intron | □ | 4 | ---- | ---- | MP |
| 22. | rs4760683 | intron | □ | 4 | ---- | ---- | MP |
| 23. | rs11168418 | intron | □ | 4 | ---- | ---- | MP |
| 24. | rs11168419 | intron | □ | 4 | ---- | ---- | MP |
| 25. | rs11168420 | intron | □ | 4 | ---- | ---- | MP |
| 26. | rs725454 | intron | □ | 5 | ---- | ---- | MP |
| 27. | rs12099462 | intron | □ | 5 | ---- | ---- | MP |
| 28. | rs879242 | intron | □ | 5 | ---- | ---- | MP |
| 29. | rs17122666 | intron | □ | 6 | ---- | ---- | MP |
| 30. | rs757556 | intron | □ | 7 | ---- | ---- | MP |
| 31. | rs12228924 | intron | ---- | 4 | ---- | ---- | NP |
| 32. | rs72644847 | intron | ---- | 4 | ---- | ---- | NP |
| 33. | rs4141065 | intron | ---- | 4 | ---- | ---- | NP |
| 34. | rs11374838 | intron | ---- | 4 | ---- | ---- | NP |
| 35. | rs10219623 | intron | ---- | 4 | ---- | ---- | NP |
| 36. | rs11168427 | intron | ---- | 4 | ---- | ---- | NP |
| 37. | rs146586156 | 5' near | ---- | 4 | ---- | ---- | NP |
| 38. | rs11168408 | 5' near | ---- | 4 | ---- | ---- | NP |
| 39. | rs10875743 | 5' near | ---- | 4 | ---- | ---- | NP |
| 40. | rs10875744 | 5' near | ---- | 4 | ---- | ---- | NP |
| 41. | rs8716 □ | syn-SNP | ---- | 5 | Tolerated | Benign | NP |
| 42. | rs72644845 | intron | ---- | 5 | ---- | ---- | NP |
| 43. | rs4760681 | intron | ---- | 5 | ---- | ---- | NP |
| 44. | rs11613781 | intron | ---- | 5 | ---- | ---- | NP |
| 45. | rs57655233 | intron | ---- | 5 | ---- | ---- | NP |
| 46. | rs7302354 | intron | ---- | 5 | ---- | ---- | NP |
| 47. | rs2107638 | intron | ---- | 5 | ---- | ---- | NP |
| 48. | rs78649179 | intron | ---- | 5 | ---- | ---- | NP |
| 49. | rs72644846 | intron | ---- | 5 | ---- | ---- | NP |
| 50. | rs11168412 | intron | ---- | 5 | ---- | ---- | NP |
| 51. | rs61343568 | intron | ---- | 5 | ---- | ---- | NP |
| 52. | rs11168421 | intron | ---- | 5 | ---- | ---- | NP |
| 53. | rs73104071 | intron | ---- | 5 | ---- | ---- | NP |
| 54. | rs58540891 | intron | ---- | 5 | ---- | ---- | NP |
| 55. | rs1859445 | intron | ---- | 5 | ---- | ---- | NP |
| 56. | rs11168413 | intron | ---- | 5 | ---- | ---- | NP |
| 57. | rs4075913 | intron | ---- | 5 | ---- | ---- | NP |
| 58. | rs11168422 | intron | ---- | 5 | ---- | ---- | NP |
| 59. | rs1476607 | intron | ---- | 5 | ---- | ---- | NP |
| 60. | rs2269933 | intron | ---- | 5 | ---- | ---- | NP |
| 61. | rs2286021 | intron | ---- | 5 | ---- | ---- | NP |
| 62. | rs80140486 | intron | ---- | 5 | ---- | ---- | NP |
| 63. | rs78294086 | intron | ---- | 5 | ---- | ---- | NP |
| 64. | rs75261947 | intron | ---- | 5 | ---- | ---- | NP |
| 65. | rs79336782 | intron | ---- | 5 | ---- | ---- | NP |
| 66. | rs75333732 | intron | ---- | 5 | ---- | ---- | NP |
| 67. | rs79751478 | intron | ---- | 5 | ---- | ---- | NP |
| 68. | rs76706444 | intron | ---- | 5 | ---- | ---- | NP |
| 69. | rs10219645 | intron | ---- | 5 | ---- | ---- | NP |
| 70. | rs10875749 | intron | ---- | 5 | ---- | ---- | NP |
| 71. | rs58556309 | intron | ---- | 5 | ---- | ---- | NP |
| 72. | rs2158515 | intron | ---- | 6 | ---- | ---- | NP |
| 73. | rs562841214 | intron | ---- | 6 | ---- | ---- | NP |
| 74. | rs34700912 | intron | ---- | 6 | ---- | ---- | NP |
| 75. | rs1049392 □ | syn-SNP | ---- | 7 | Tolerated | Benign | NP |
| 76. | rs58331734 | intron | ---- | 7 | ---- | ---- | NP |
| 77. | rs534307036 | intron | ---- | 7 | ---- | ---- | NP |
| 78. | rs113785514 | intron | ---- | 7 | ---- | ---- | NP |
| 79. | rs527761146 | intron | ---- | 7 | ---- | ---- | NP |
| 80. | rs10875747 | intron | ---- | 7 | ---- | ---- | NP |
| 81. | rs10875748 | intron | ---- | 7 | ---- | ---- | NP |
| 82. | rs11168424 | intron | ---- | 7 | ---- | ---- | NP |
| 83. | rs77966213 | intron | ---- | 7 | ---- | ---- | NP |
| 84. | rs11168425 | intron | ---- | 7 | ---- | ---- | NP |
| 85. | rs148905596 | intron | ---- | 7 | ---- | ---- | NP |
| □ denote conserved SNPs, ✓ SNPs which affect function, ---- SNPs with no detected function, NF = SNP not found,<br>HP = Highly prioritized (Detected functional by two or more tools)<br>MP = Moderately prioritized (Detected functional by one tool only)<br>NP = Not/Poorly prioritized (Not found to be functional by any tool) |  |  |  |  |  |  |  |

## DISCUSSION

Elevated aerobic glycolysis, a characteristic of cancer cells (Warburg effect) is energetically inefficient, that triggers cachexia (irreversible muscle wasting) in cancer patients leading to death [31–32]. This wasting syndrome reduces drug effectiveness in patients fighting against cancer. The prevalence of muscle loss varies with the type of cancer (*e.g*. 16% in breast cancer), stage of cancer and host’s response to cancer [33]. Therefore, abnormalities in metabolic genes particularly those encoding glycolytic regulatory proteins can be targeted as a potential therapeutic intervention strategies for the killing cancer cells and coping co-morbid conditions. For this, one such probable therapy can be inhibition and/or down regulation of human PFKM to lowers the survival of cancer cells. PFKM is an imperative glycolytic regulatory target for the reason that it serves as energetic activator of muscle glycolysis, which is crucial for the dissemination of the cancer [34].

It’s the genes that codes for the formation of functional proteins. Therefore, identification of genetic variations that interfere in the expression and/or structure of PFKM that subsequently alter protein structure/function will be clinically important. The commonly found genetic variations is SNPs, as for instance human *PFKM* consists total of 9694 SNPs *per se* (source dbSNP database, dated October 2019). For this, selection of variants could be made on the basis of putative functional effects *e.g*. altering transcription, translation or miRNA binding from the SNPs that may not have any functions or are rare. As the interpretation of clinically important novel variants often remain challenging, many bioinformatics tools have been developed that predict biological consequences of these polymorphisms. Therefore, an effort was made to haul out the most likely functional variants of human *PFKM* using *in silico* analysis, so that the focus can be put on describing those SNPs which probably may have an important role in controlling the disease condition.

However, no single bioinformatics tool can completely depict the functional significance of allelic variants. Hence, the current analysis was conducted using a number of complementary bioinformatics tools, for instance, Ensembl Genome browser was used for the screening of conserved SNPs (cSNPs), Two composite tools [FuncPred (SNPinfo), and RegulomeDB (v 2.0)] were used the function prediction, SIFT and PolyPhen-2 were used to predict the phenotypic effect of coding SNPs, on the physio-chemical properties of *PFKM* and thus its function. Different algorithms for different softwares can sometime leads to the mis-interpretation of biological data accredited to non-overlapping functions prediction. To evade such discrepancies, two major parameters were considered before commencing *in silico* functional predictions. These parameters includes, verifying the validation status of a given SNP to avoid inclusion of fabricated ones [35]. Secondly, use of MAF cut-off of ≥0.10 that denotes its 10% frequency in a given population, which is ultimately crucial for SNP study in relation to disease with lower prevalence [36]. Considering these factors and outcomes of softwares employed, the present study filtered 11 SNPs of *PFKM* that has highest likelihood of being functionally important. These 11 SNPs include both non-coding (rs11168417, rs4760619, rs12306290, rs11168416, rs10492080, rs11168415, rs10875745, rs2269935 and rs10082861) as well as coding (rs11609399 and rs2228500) SNPs. Of the prioritized SNPs, most variants did not have any previous reports in relation to cancer, while the role of some SNPs in cancer etiology has been previously reported, which was used as an evidence to authenticate the findings of the present study.

The non-coding variants were found to be altering transcription factor binding sites (TFBS), resultantly affecting the gene expression. A study by Konieczna *et al*. [37] performed the partial silencing of *PFKM* experimentally and observed down-regulation of *PFKM* expression leading to impaired DNA synthesis and thus subsequent arrest of cells in G1 phase. The study further suggested genetic polymorphisms in *PFKM* can leads to the same results causing alteration in cells metabolism *via* inhibition of cell cycle and cellular apoptosis, an essential property of tumor cells. Apart from this, studies have investigated the effect of these polymorphisms in relation to different diseases. HP-SNP rs2269935 of *PFKM* was found to be associated with relative lowering in body fat [38]. Moreover, the HP variant of *PFKM i.e*. rs11168417 has been depicted as a prognostic marker for non-small cell lung cancer in humans undergoing surgical resection/chemotherapy [4, 39]. The studies suggested the important role of this polymorphism in determining the prognosis of patients with lung cancer as it affects macromolecular biosynthesis, energy production, and other non-glycolytic functions in cancer cells.

Of the HP-coding variants, ns-SNP rs11609399 was found to be deleterious accredited to amino-acid conversion from non-polar, Leucine to positively charged, Histidine at ammino-acid position ‘2’ of protein, thus affecting protein structure and function. Moreover, both HP-coding (rs11609399 and rs2228500) variants were found to be evolutionary conserved and affecting exon splicing. The cSNPs are more likely to have biological functions [40], while aberrant splicing by these variants may abrogate PFKM synthesis/function, and thus can effect cancer susceptibility. In consonance to this, gene based analysis of *PFKM* by Ahsan *et al*. (2014) [18] suggested *PFKM* serves as a hub of genetic variations including rs2228500, and thus serve as susceptibility locus for early onset of breast cancer (EOBC) in Caucasian women of all ages. Since *PFKM* is a glycolytic enzyme, which regulates aerobic glycolysis in cancer cells, it can be hypothesized that genetic alterations detected in human PFKM may contribute to other cancer types as well.

Thus, the prioritized putative functional SNPs may have profound biological significance in relation to cancer for several reasons. The foremost reason is the expression of *PFKM* in cancer cell lines [41]. Secondly, *PFKM* variants have been linked with post-translational modifications resulting in dissemination of cancer cells and altered metabolism [11]. Thirdly, an association between *PFKM* and cancer risk is reasonable due Warburg effect being employed by cancer cells for performing high rate of glucose metabolism *via* aerobic glycolysis [11]. Finally, p53 is known to prevent the embryo development in model organism with defects/mutations in genes including *PFKM* [42]. Since the science behind PFKM and its regulators is well described [43–44], establishment of *PFKM* as a cancer gene and its highly ranked functional SNPs as genetic markers may provide potential translational implications for cancer prevention and treatment.

In **conclusion**, PFKM plays a pivotal role in feeding cancer cells *via* Warburg phenomena. Therefore, an effort was made to identify its casual variants that may affect protein function and cancer possibility. The study identified 11 *PFKM* SNPs including rs11609399, rs11168417, rs4760619, rs12306290, rs11168416, rs10492080, rs11168415, rs10875745, rs2269935, rs10082861and rs2228500 as putative functional variants, which is worthy of further investigation to confirm their predictive role in cancer etiology. Once confirmed, these SNPs can serve as genetic markers for cancer prevention, prognosis, diagnosis and treatment strategies. Hence, the present research serve as base for future studies associated with glycolytic genes in cancers.

## Author’s contributions

YR reviewed the literature, was involved in design, performing analysis, interpretation, and drafted the manuscript. KK and MS contributed manuscript editing. NK supervised the research, reviewed the literature, designed the experiment, data analysis, manuscript editing and supervision. All authors read and approved the final manuscript.

## Availability of data and materials

The data that support the findings of this study are available as supplementary material. Additional information can be obtained from the corresponding author upon reasonable request.

## Consent for publication

All authors give their consent for manuscript publication.

## Acknowledgements

Not applicable

## Ethics approval and consent to participate

Not applicable

