## Supplementry Material for "*In silico* analysis of SNPs in human phosphofructokinase, Muscle (*PFKM*) gene: An apparent therapeutic target of aerobic glycolysis and cancer"

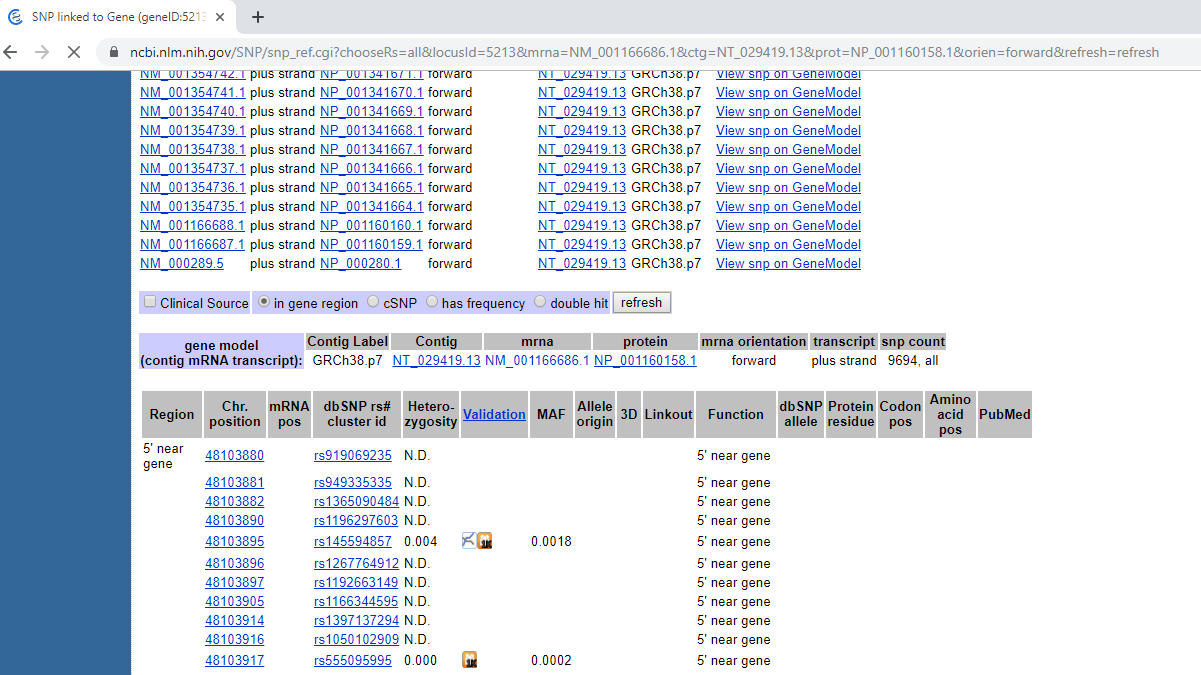

**S. Fig. 1:** Image showing the SNP gene-view region of human *PFKM* gene in NCBI (Screenshot view).

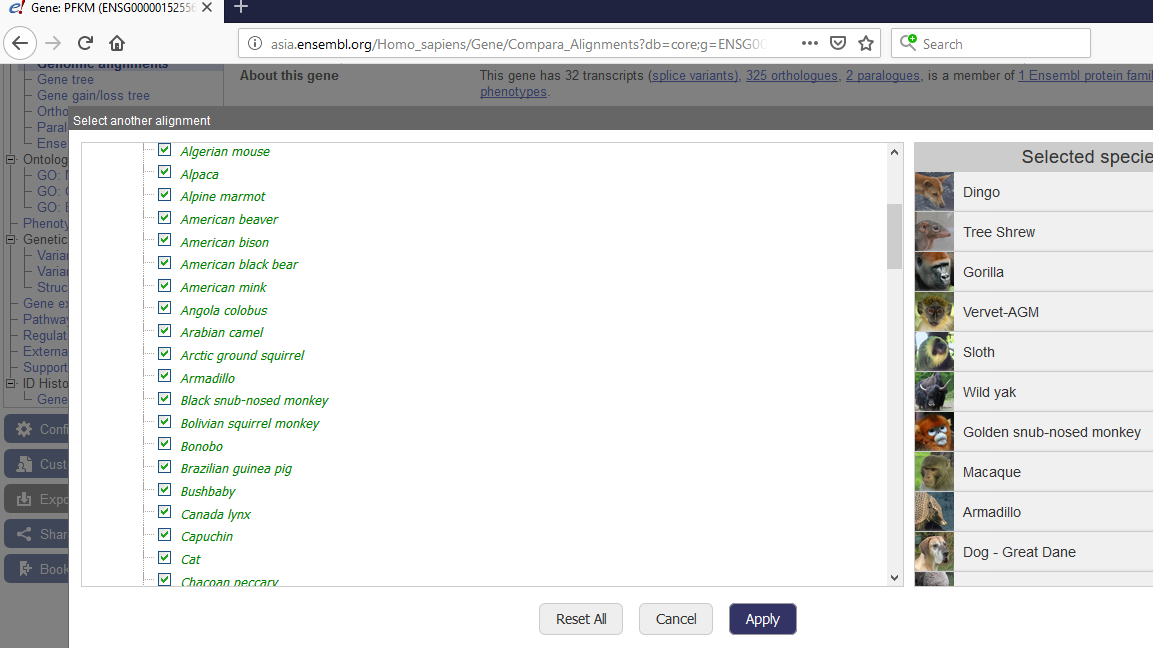

**S. Fig. 2:** Image showing some species from 91 eutherian mammals species selected from genomic alignments, this screenshot was taken from Ensembl Genome browser 98 (Screenshot view).

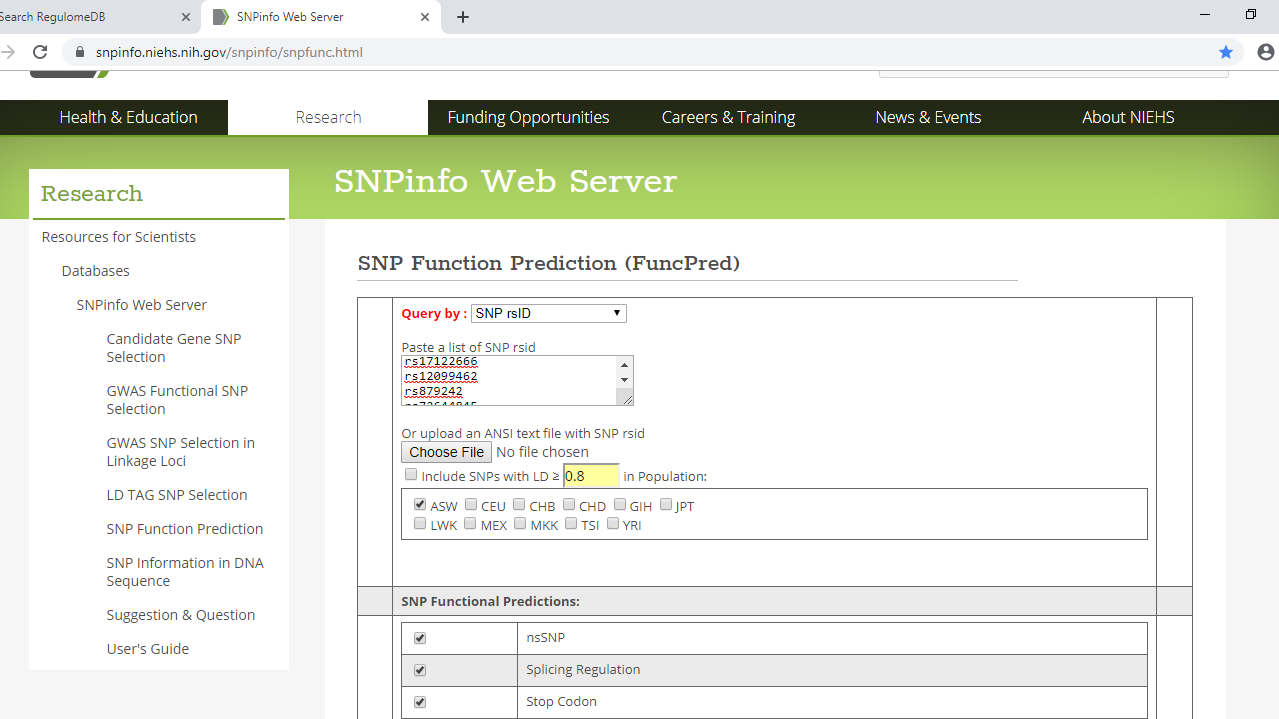

**S. Fig. 3:** The image showing scheme of data input in the form of rsIDs list of validated SNPs in FuncPred tool with default setting in Asian population (ASW), this screenshot was taken from SNPinfo (FuncPred) bioinformatics tool (Screenshot view).

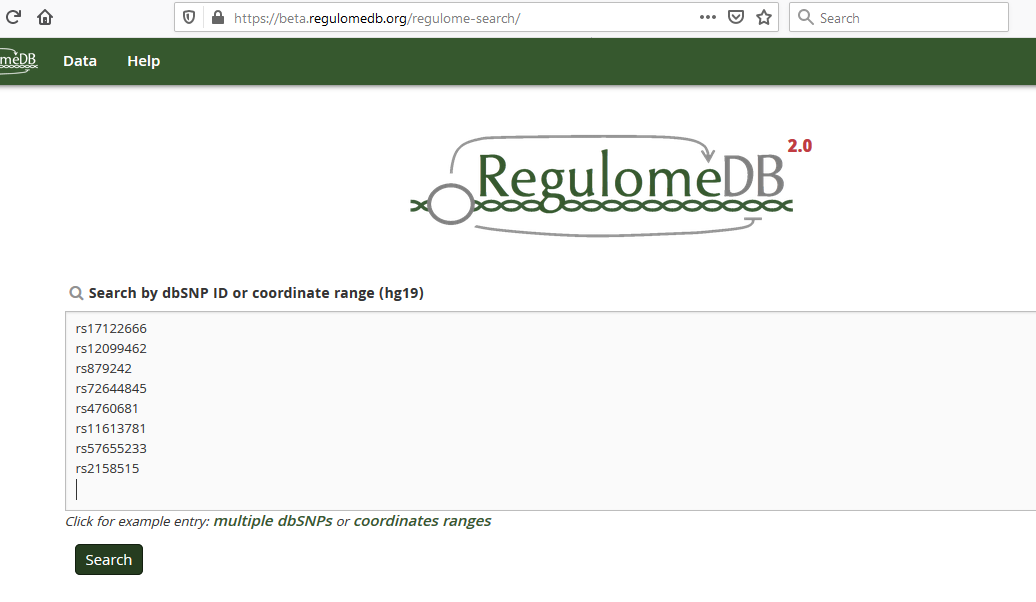

**S. Fig. 4:** The image showing scheme of data input in the form of rsIDs of validated SNPs with default settings, this screenshot was taken from Beta Regulome database (Screenshot view).

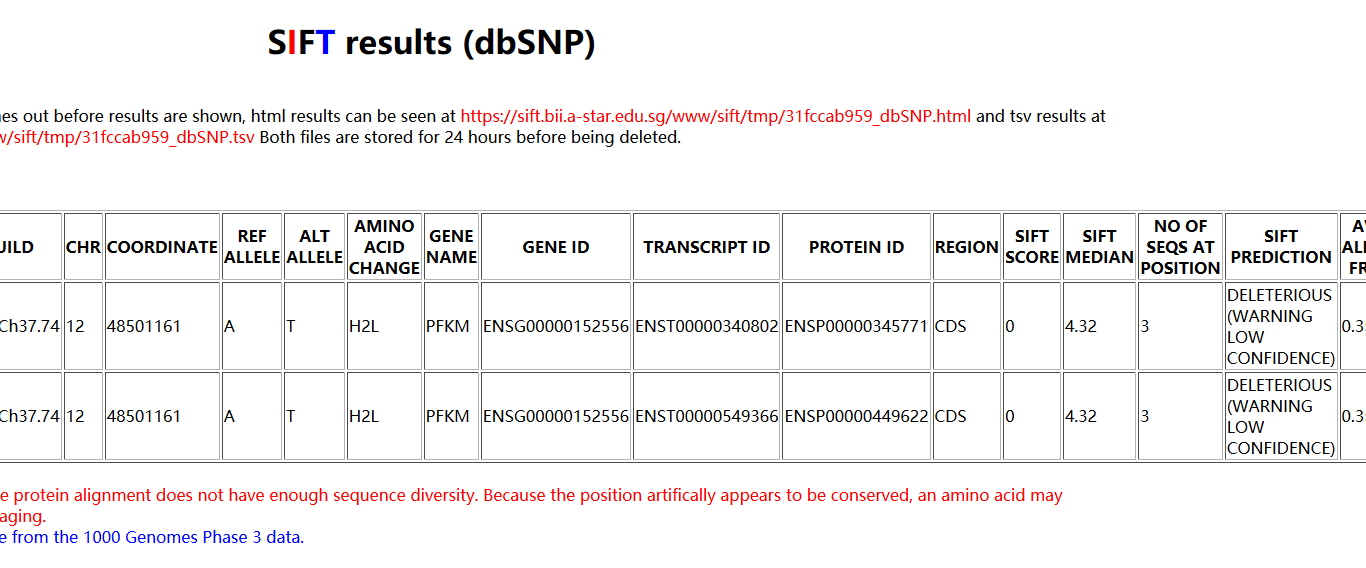

**S. Fig. 5:** Image showing the prediction of nsSNP rs11609399 of human *PFKM* in SIFT tool having deleterious effect (Screenshot view).

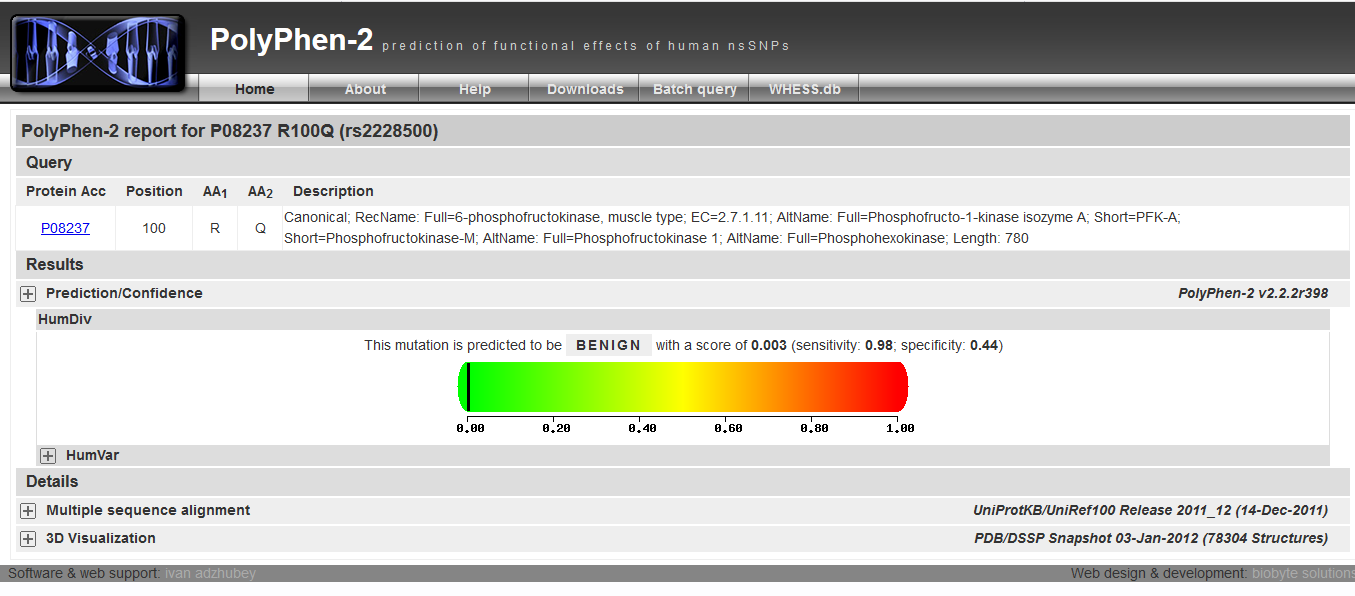

**S. Fig. 6:** Image showing the prediction outcome of benign effect of nsSNP rs2228500 with score of 0.003,(Screenshot view).

**S. Table 1** – Beta Regulome database annotation scores representing the category with description.

| CATEGORY | | DESCRIPTION |
| --- | --- | --- |
| Likely to affect binding and linked to expression of a gene target | | |
| 1b | eQTL + TF binding + any motif + DNase footprint + DNase peak | |
| 1c | eQTL + TF binding + Matched TF motif + DNase peak | |
| 1d | eQTL + TF binding + Any motif + DNase peak | |
| 1e | eQTL + TF binding + Matched TF motif | |
| 1f | eQTL + TF binding/DNase peak | |
| Likely to affect binding | | |
| 2a | TF binding + Matched TF motif + Matched DNase footprint + DNase peak | |
| 2b | TF binding + any motif + DNase footprint + DNase peak | |
| 2c | TF binding + Matched TF motif + DNase peak | |
| Less likely to affect binding | | |
| 3a | TF binding + any motif + DNase peak | |
| 3b | TF binding + Matched TF motrif | |
| Minimal binding evidence | | |
| 4 | TF binding + DNase peak | |
| 5 | TF binding or DNase peak | |
| 6 | Motif hit | |
